# Muscle Moment Arm–Joint Angle Relations in the Hip, Knee, and Ankle: A Visualization of Datasets

**DOI:** 10.1101/2024.06.22.600198

**Authors:** Ziyu Chen, David W. Franklin

## Abstract

Muscle moment arm is a property that associates muscle force with joint moment and is crucial to biomechanical analysis. In musculoskeletal simulations, the accuracy of moment arm is as important as that of muscle force, and moment arm calibration requires data from anatomical measurements. Nonetheless, such data are elusive, and the complex relation between moment arm and joint angle can be unclear. Using common techniques in systematic review, we collected a total of 300 moment arm datasets from literature and visualized the muscle moment arm–joint angle relations in the hip, knee, and angle. The findings contribute to the understanding and analysis of musculoskeletal mechanics, as well as providing reference regarding the experimental design for future moment arm measurements.

## 1. Introduction

The movement of the human body is in essence the compound motion of joints actuated by torques. More precisely, it is driven by *moments of force*, since the actuation originates from forces exerted by muscles, which are not force couples such as what generally arise from electric motors (Kane and Levinson, 1985; Yamaguchi, 2001). Thus, an important difference between the studies of robotics and human kinetics is that biomechanists must take into account the property that transforms muscle force into joint moment, namely muscle moment arm (van den Bogert et al., 2013; Pandy, 1999; Zajac, 1993). The sign of moment arm determines the role of the muscle to a joint, e.g., whether it is an extensor or flexor, and its magnitude partially determines the level of muscle activation required in a motion.

In simple cases, moment arm can be perceived as the distance between the muscle line of action and the center of rotation (CoR), and many experiments have utilized this idea to measure moment arm (Hashizume et al., 2012; Herzog and Read, 1993; Maganaris et al., 2000; Rugg et al., 1990; Smidt, 1973). However, the path of a muscle is often not straight, and the distance between its *curve* of action and the rotation center is no longer a constant value, even when the muscle is isometric and at rest. In such scenario, the CoR method will be inaccurate and one must visit the precise definition of moment arm for solutions.

In strict terms, joint moment is associated with muscle force by:

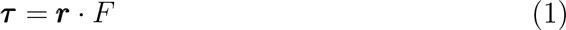

where *τ* is an *N* × 1 vector of moment, *F* is a scalar of force, and *r* is an *N* × 1 vector of moment arm; *N* is the number of degrees of freedom (DoF) in the system.

Hence, if both joint moment and muscle force are known, moment arm is easily calculated as their quotient. However, although joint moment is simple to measure, muscle force is difficult to obtain *in vivo*, and this concept of *kinetic balance* (KB) is only occasionally employed in cadaveric studies (Aper et al., 1996; Weinstein et al., 1987).

A practical alternative is to follow the principle of virtual work (Sherman et al., 2013): Since muscle force is transformed into joint moment, then in any instant, the work done by the muscle should theoretically equal to the work done by the joint:

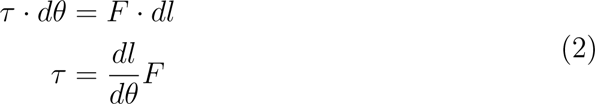

where *dθ* and *dl* are small displacements in joint angle and muscle length.

Then, from Eq 1 we may have:

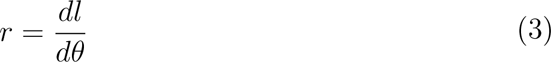

which serves as an alternative definition for moment arm. With this, moment arm can be calculated based on the changes in joint angle and muscle length, the latter of which is much easier to measure *in vivo* than muscle force. This is the tendon excursion (TE) method and frequently used for moment arm measurement (Eng et al., 2015; Hintermann et al., 1994; Maganaris et al., 2000; McCullough et al., 2011; Spoor et al., 1990).

One other definition of moment arm surfacing in literature (Gray et al., 2021; Krevolin et al., 2004; Pandy, 1999) is:

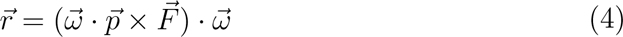

where 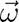 denotes the axial direction of joint rotation in a 3D Cartesian space, 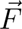 is a 3D unit vector denoting the direction of the muscle line of action, and 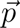 is a 3D position vector from any point on the rotation axis to the muscle line of action. Note that 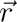 is a 3D vector of moment arm, different from ***r*** in Eq 1; specifically, the 2-norm of 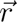 could be one of the elements in the vector ***r***.

Critically, this definition assumes the existence of the line of action. Therefore, for application, one must decide which part of the muscle is treated as the straight path to yield 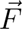 for moment arm calculation, and this is es-sentially still the CoR method.

With the relation between kinetics and moment arm elucidated, it is clear how crucial moment arm is to biomechanical analysis. For example, for accu-rate musculoskeletal simulation, there has been a major focus on contraction dynamics, such as improving muscle models (Millard et al., 2013, 2023), investigating the sensitivity of muscle force estimation to musculotendon pa-rameters (Blache et al., 2019; Chen and Franklin, 2023; Claiborne et al., 2006), and performing large-scale human measurements for model calibra-tion (Handsfield et al., 2014; Ward et al., 2009). Nevertheless, these efforts contribute solely to the accuracy of *F* in Eq 1, while ***r*** is equally crucial in predicting moment output or estimating muscle activation. It is thus impor-tant to know how moment arm changes with different joint positions, as well as which muscles lack sufficient measurement of moment arm.

The purpose of this study is to collect existing datasets from literature constituting moment arm–joint angle relations in the hip, knee, and ankle. We approach the issue using common techniques in systematic review and we aim to visualize data with sufficient details to provide reference values for biomechanical analysis, especially musculoskeletal modeling.

## 2. Methods

Moment arm in the lower limb is investigated by muscle group. For each muscle group, we used a combination of keywords to initiate batch searches in Google Scholar with the software Publish or Perish. To guarantee search efficiency, we experimented multiple combinations of keywords in search of 17 target studies with Achilles tendon moment arm data and 9 studies with patellar tendon moment arm data. The keyword set selected for the final batch search is one that led to the most target papers while having them appear in top search rankings: For the Achilles tendon, it was 13 out of 17 and on average 11^th^ place in the list of 200 results, while it was 9 out of 9 and 18^th^ place for the patellar tendon.

The format of this keyword set is shown in Table 1, where the number of searches for each muscle group is based on muscle size (Handsfield et al., 2014). We presume it is easier to measure moment arm on large muscles, so there should be more studies reporting relevant data, and more searches are required for larger muscles. In total, 4400 searches were initiated for 21 muscles or muscle groups.

**Table 1.** Search number and keywords for each muscle or muscle group.

| Search Number | Keyword Set (Part 1)* |
| --- | --- |
| 500 | (triceps surae OR achilles)<br>(quadriceps OR patella)<br>gluteus<br>(hamstring OR biceps femoris<br>OR semimembranosus OR semitendinosus) |
| 200 | (iliopsoas OR iliacus OR psoas)<br>tibialis posterior<br>adductor AND (magnus OR brevis OR longus)<br>flexor AND (hallucis OR digitorum longus)<br>peroneus<br>tibialis anterior<br>extensor AND (hallucis OR digitorum longus) |
| 100 | pectineus<br>piriformis<br>gracilis<br>tensor fascia<br>sartorius<br>obturator<br>popliteus<br>gemelli OR gemellus<br>quadratus femoris<br>plantaris |

\*Part 2: AND moment arm AND (rotation OR excursion).

Other details of data collection are shown in Fig 1 and can also be found in Appendix A. Note that we also initiated 4400 searches oriented towards joint moment for another study, but the search results were examined altogether. This serves as a compensation for the potential overfitting of the keywords from Table 1, in case some records are missed in the searches intended for moment arm data.

**Figure 1.**
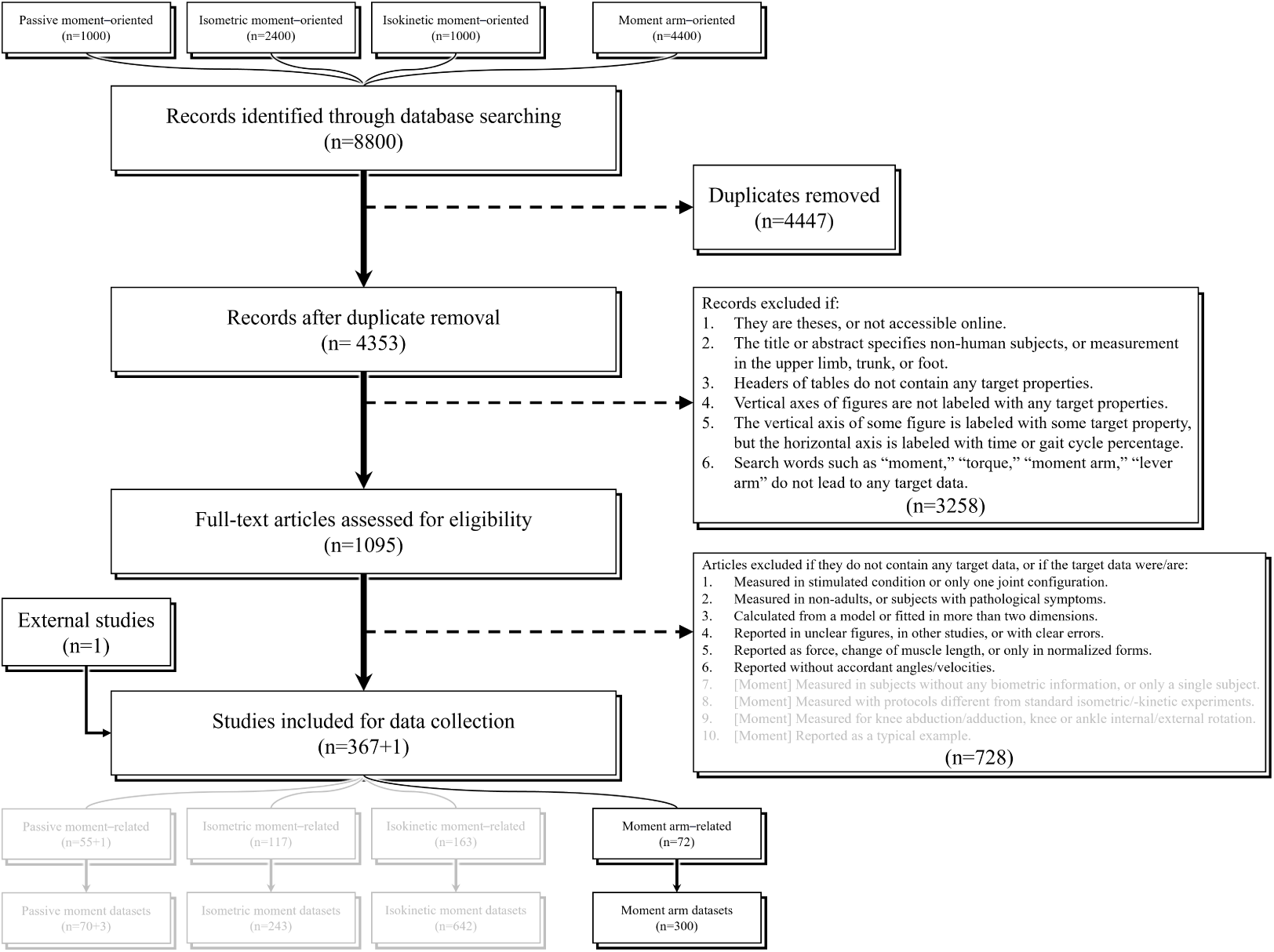
Workflow of study selection. Moment–related datasets are not presented in this study, but records identified by searches intended for moment data were examined for moment arm data.

In data collection, if a study presents its data in graphs rather than in numbers, we use Graph Grabber for digitization. The data are categorized by attributes listed in Table 2, and a dataset is generally defined as a set of measurements with distinction in either its reference, subject information, measured muscle, primary DoF, or data type. A muscle may have multiple moment arms, and the measured moment arm is indicated by the primary DoF. Meanwhile, data type denotes the method of measurement:

1. Two-dimensional center of rotation (CoR)
2. Three-dimensional center of rotation (CoR3)
3. Tendon excursion (TE)
4. Kinetic balance (KB)

**Table 2.**
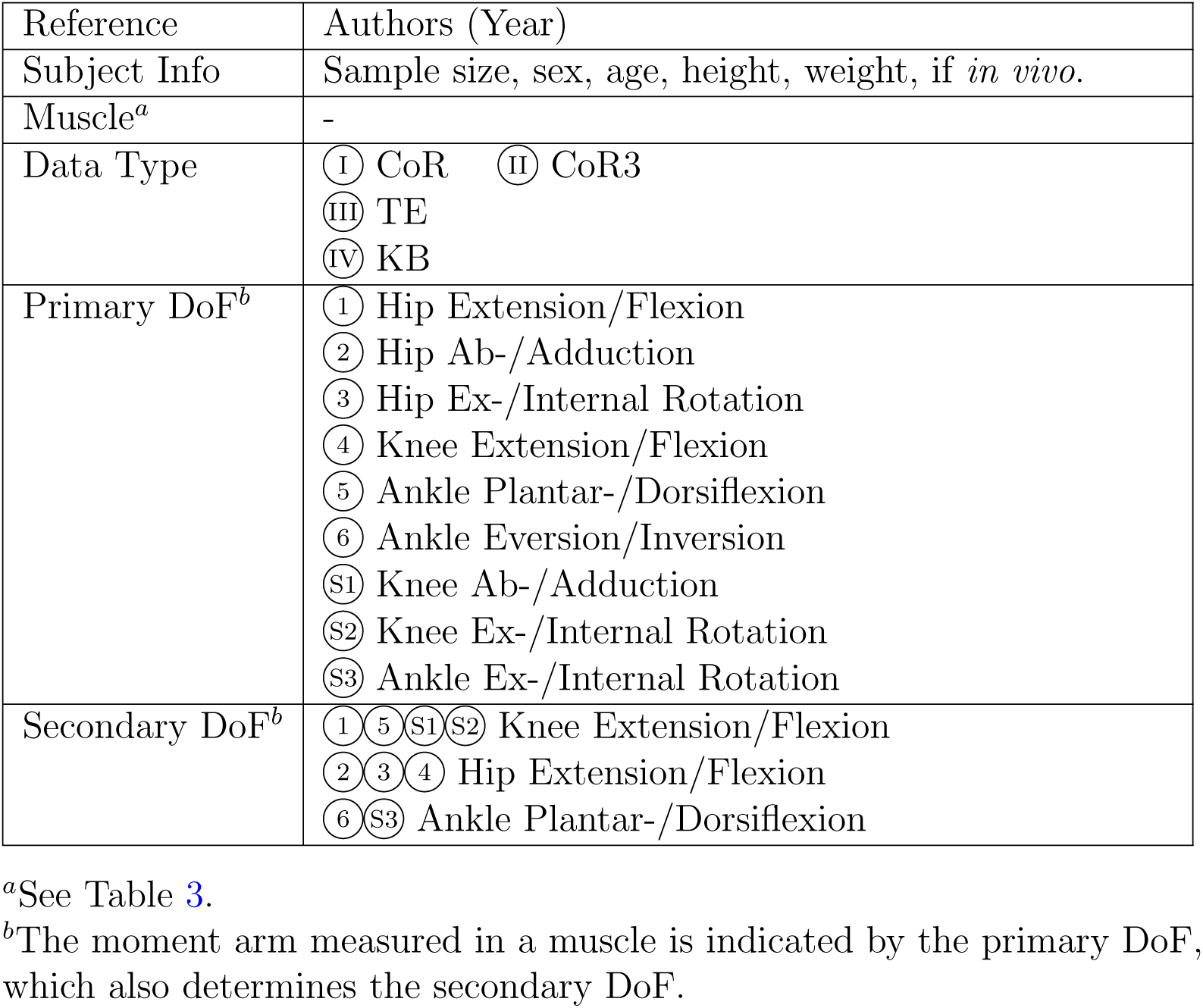
Metadata for moment arm datasets.

When measured *in vivo*, a muscle could be in different states of activation, and for the potential need of comparative analysis, we labeled the data type with the plus sign as an approximate indication of muscle activation: not activated (no sign), activated up to 33% (+), 34%–67%(++), and 67%–100%(+ + +). In some studies, a muscle is treated as multiple strings or measured with the same method but different configurations (e.g., CoR defined as different anatomical landmarks), and the results are recorded as different datasets with the configurations noted. For convenience, we provide a catalog with details of the datasets (Appendix A).

In each dataset, there is at least one curve, which is comprised of measurements from at least two joint positions, and some datasets contain multiple curves, each comprised of multiple measurements. The data for each curve are stored as a 3-column matrix, with the first, second, and third column for the angles in the primary and secondary DoFs, and the moment arms. The sign rule for angle and moment arm in each DoF is based on the ISB recommendations (Wu et al., 2002; Derrick et al., 2020) except for knee flexion/extension, which is reversed so that moment arms for anti-gravity motions share the negative sign. If the angle in the secondary DoF is not specified in a study, we label it as NaN.

Since the data are mostly two dimensional and the measured joint positions are different across datasets, typical methods of meta-analysis no longer apply, and we did not proceed to combine the results. Also, considering the heterogeneity between studies, we refrain from making conclusive statements for any moment arm on the representative magnitude. For biomechanical analysis, readers are encouraged to select datasets with characteristics of subjects and means of measurement conforming to their object of study.

## 3. Results

As shown in Fig 1, a total of 72 studies were identified from 4353 records searched in the database, and they yielded 300 moment arm datasets. Table 3 shows the distribution of datasets. See Appendix B for a summary of collected studies and datasets.

**Table 3.** Distribution of datasets for moment arms in each lower limb muscle.

| Muscle | Number of Datasets |  |  |  |  |  |  |  |  |
| --- | --- | --- | --- | --- | --- | --- | --- | --- | --- |
|  | Hip |  |  | Knee |  |  | Ankle |  |  |
|  | S | F | T | S | F | T | S | F | T |
| Achilles tendon |  |  |  |  |  |  | 38 | 3 | 1 |
| Soleus |  |  |  |  |  |  | 1 | - | - |
| Gastrocnemii |  |  |  | 1 | - | - | - | - | - |
| Gastrocnemius med. |  |  |  | 5 | - | - | 2 | 3 | - |
| Gastrocnemius lat. |  |  |  | 5 | - | - | 1 | 3 | - |
| Patellar tendon |  |  |  | 40 | - | - |  |  |  |
| Quadriceps tendon |  |  |  | 4 | - | - |  |  |  |
| Rectus femoris | 1 | - | - | 7 | - | - |  |  |  |
| Vastus medialis |  |  |  | 4 | - | - |  |  |  |
| Vastus lateralis |  |  |  | 4 | - | - |  |  |  |
| Vastus intermedius |  |  |  | 3 | - | - |  |  |  |
| Gluteus maximus | 4 | 4 | 6 | 4 |  |  |  |  |  |
| Gluteus medius | 3 | - | 6 |  |  |  |  |  |  |
| Gluteus minimus | - | - | - |  |  |  |  |  |  |
| Biceps femoris |  |  |  | 6 | - | - |  |  |  |
| short head |  |  |  | 1 | - | 1 |  |  |  |
| long head | 1 | - | - | 4 | - | 1 |  |  |  |
| Semimembranosus | 1 | - | - | 8 | - | 1 |  |  |  |
| Semitendinosus | 1 | - | - | 9 | - | 1 |  |  |  |
| Iliopsoas | - | - | 1 |  |  |  |  |  |  |
| Psoas major | 1 | - | - |  |  |  |  |  |  |
| Iliacus | - | - | - |  |  |  |  |  |  |
| Tibialis posterior |  |  |  |  |  |  | 5 | 7 | 1 |
| Adductor lon./bre./mag. | - | - | - |  |  |  |  |  |  |
| Flexor hallucis longus |  |  |  |  |  |  | 4 | 3 | 1 |
| Flexor digitorum longus |  |  |  |  |  |  | 2 | 3 | 1 |
| Peroneus longus |  |  |  |  |  |  | 3 | 3 | 1 |
| Peroneus brevis |  |  |  |  |  |  | 3 | 3 | 1 |
| Tibialis anterior |  |  |  |  |  |  | 16 | 8 | 1 |
| Extensor hallucis longus |  |  |  |  |  |  | 2 | 2 | 1 |
| Extensor digitorum longus |  |  |  |  |  |  | 2 | 2 | 1 |
| Pectineus | - | - | - |  |  |  |  |  |  |
| Piriformis | 1 | 1 | 1 |  |  |  |  |  |  |

| Muscle | Number of Datasets |  |  |  |  |  |  |  |  |
| --- | --- | --- | --- | --- | --- | --- | --- | --- | --- |
|  | Hip |  |  | Knee |  |  | Ankle |  |  |
|  | S | F | T | S | F | T | S | F | T |
| Gracilis | - | - | - | 7 | - | 1 |  |  |  |
| Tensor fasciae latae | 2 | 2 | - | 3 |  |  |  |  |  |
| Sartorius | - | - | - | 5 | - | 1 |  |  |  |
| Obturator internus | 1 | 1 | 2 |  |  |  |  |  |  |
| Obturator externus | 2 | - | 1 |  |  |  |  |  |  |
| Popliteus |  |  |  | 1 | - | 1 |  |  |  |
| Gemelli | - | - | - |  |  |  |  |  |  |
| Quadratus femoris | 2 | - | 1 |  |  |  |  |  |  |
| Plantaris |  |  |  | - | - | - | - | - | - |

S: sagittal motion (hip or knee extension/flexion, ankle plantar-/dorsiflexion). F: frontal motion (hip or knee ab-/adduction, ankle eversion/inversion). T: transverse motion (ex-/internal rotation). The dash indicates that the muscle potentially actuates this DoF but no study has reported the accordant moment arm.

The gathered datasets are mostly visualized in Figs 2–10. The sex of the subject group is indicated by the color of the scatters: red (predominantly female), blue (predominantly male), purple (mixed or unknown). The circle shape indicates *in vivo* measurements on young subjects, whereas the cross mark if measurements are from cadeveric specimens. The method of mea-surement is denoted by the line style: solid (CoR), dashed (TE), dash-dotted (KB). Only two datasets were measured using on the KB method (Aper et al., 1996; Weinstein et al., 1987), whereas 189 were from the TE method. The CoR method contributed 109 datasets, of which 24 were measured based on a 3D center.

**Figure 2.**
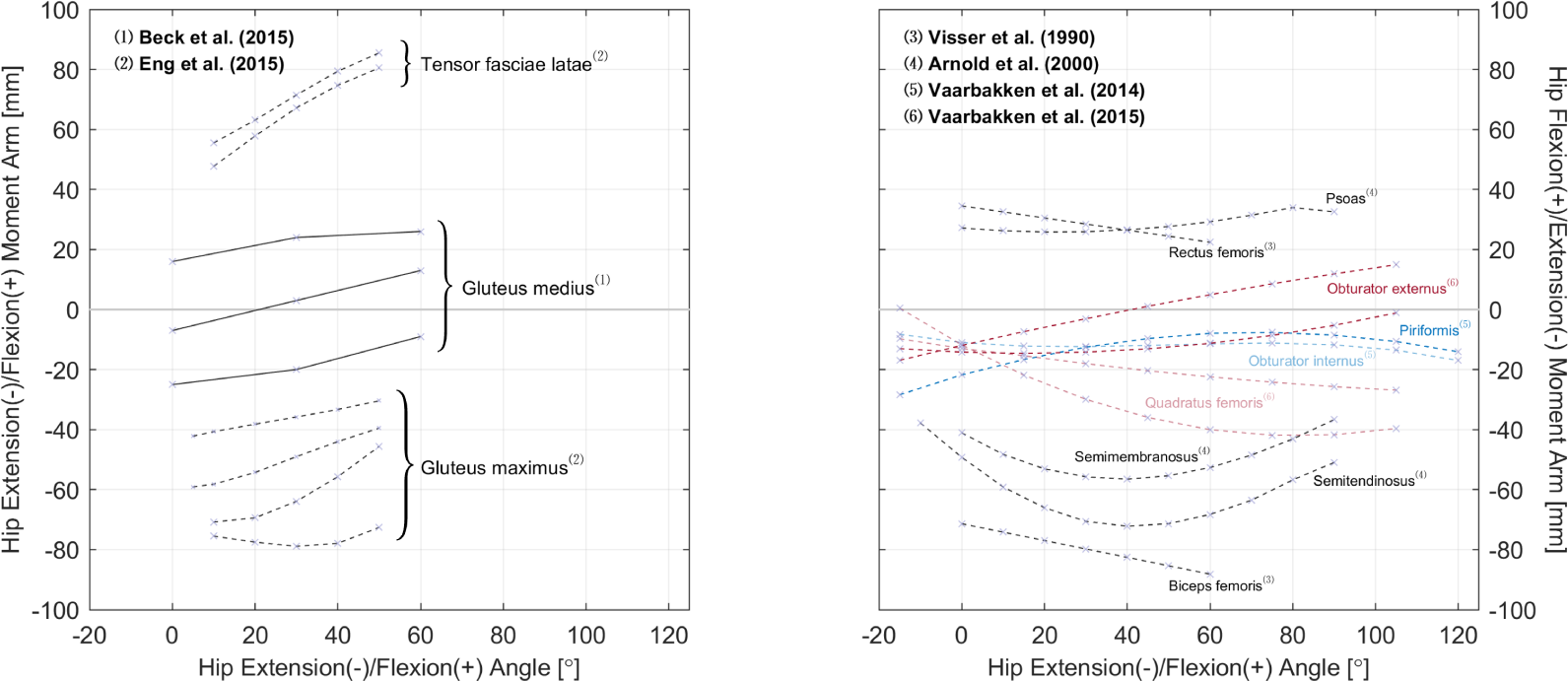
Relations of hip extension/flexion moment arms. Left: the gluteus maximus, gluteus medius, and tensor fasciae latae. Right: the rectus femoris and other gluteal muscles with available data. All measurements were performed on cadeveric specimens of mixed/unknown sex (purple cross mark). The solid and dashed line styles denote respectively the CoR and TE methods; the different line and text colors are merely for convenience of distinction.

**Figure 3.**
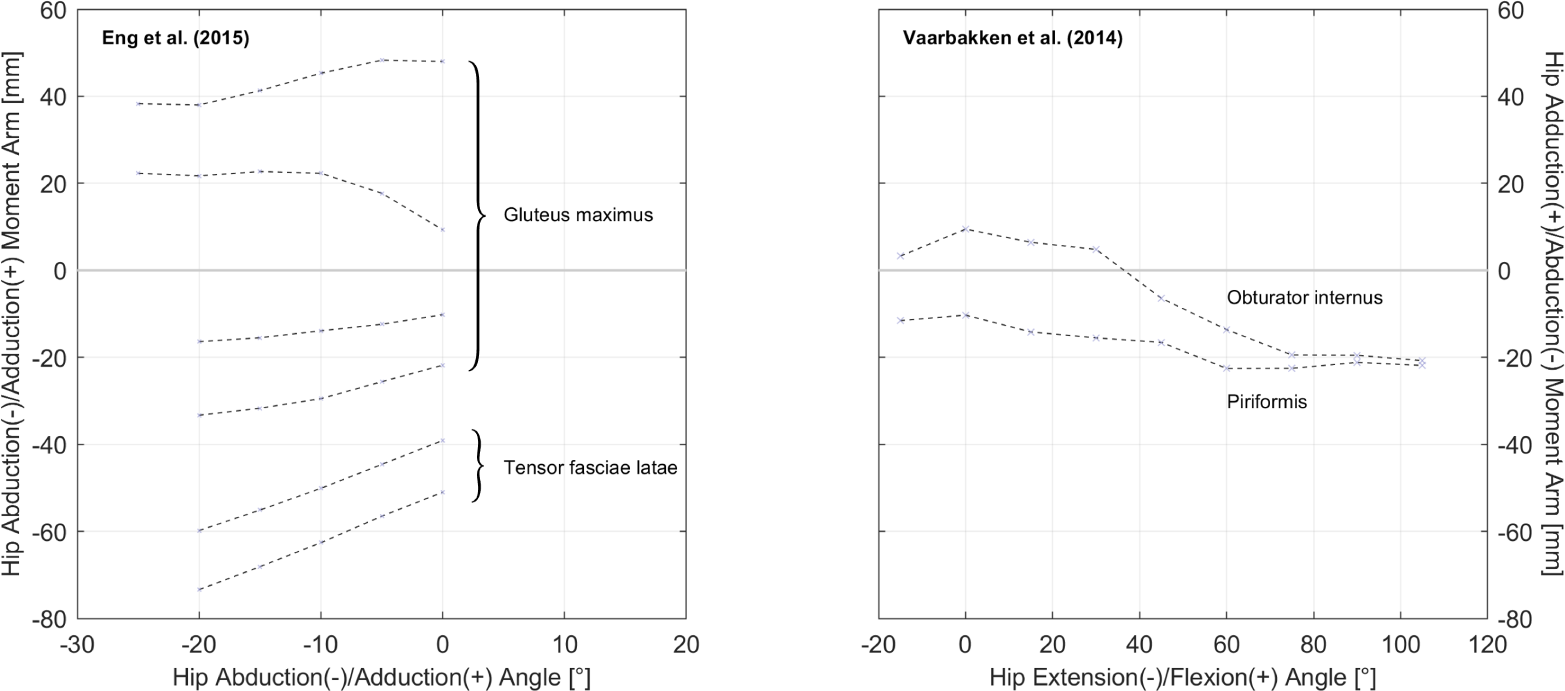
Relations of hip ab-/adduction moment arms. Left: relations with hip adduction angle. Right: relations with hip flexion angle. All measurements were performed on cadeveric specimens of mixed/unknown sex (purple cross mark) with the TE method (dashed line style).

**Figure 4.**
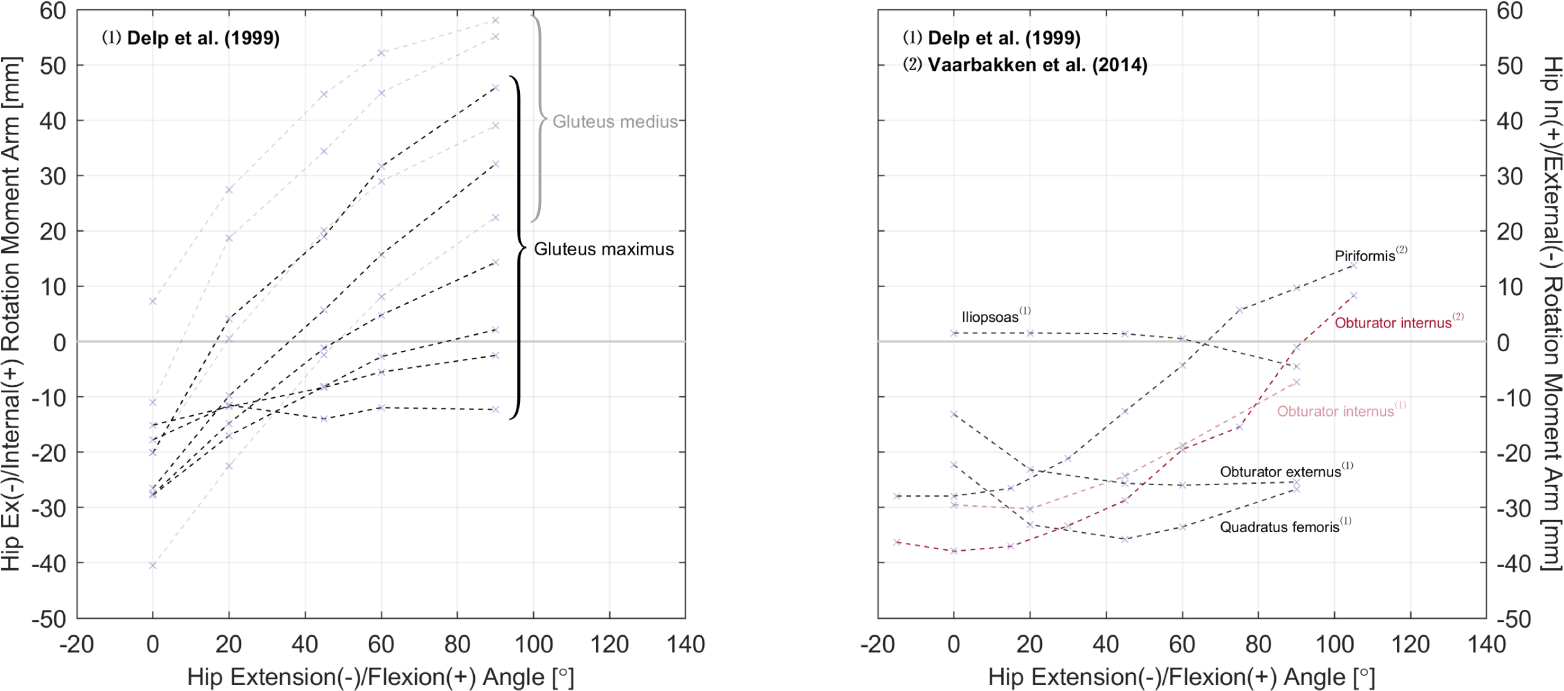
Relations of hip ex-/internal moment arms. Left: the gluteus maximus and medius. Right: other gluteal muscles with available data. All measurements were performed on cadeveric specimens of mixed/unknown sex (purple cross mark) using the TE method (dashed line style); the different line and text colors are merely for convenience of distinction.

**Figure 5.**
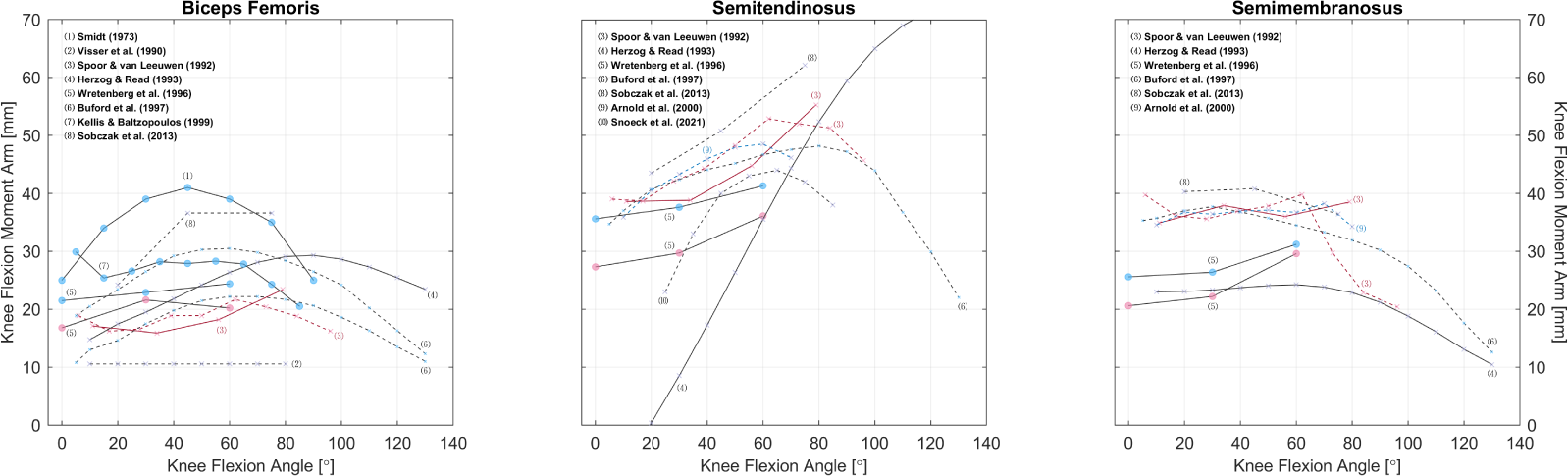
Relations of knee flexion moment arms of the hamstrings. Measurements were performed on healthy subjects (circle) or cadeveric specimens (cross mark). The scatter colors of red, blue, and purple denote respectively the sex of male, female, and mixed/unknown. The solid and dashed line styles denote respectively the CoR and TE methods; the different line and text colors are merely for convenience of distinction.

**Figure 6.**
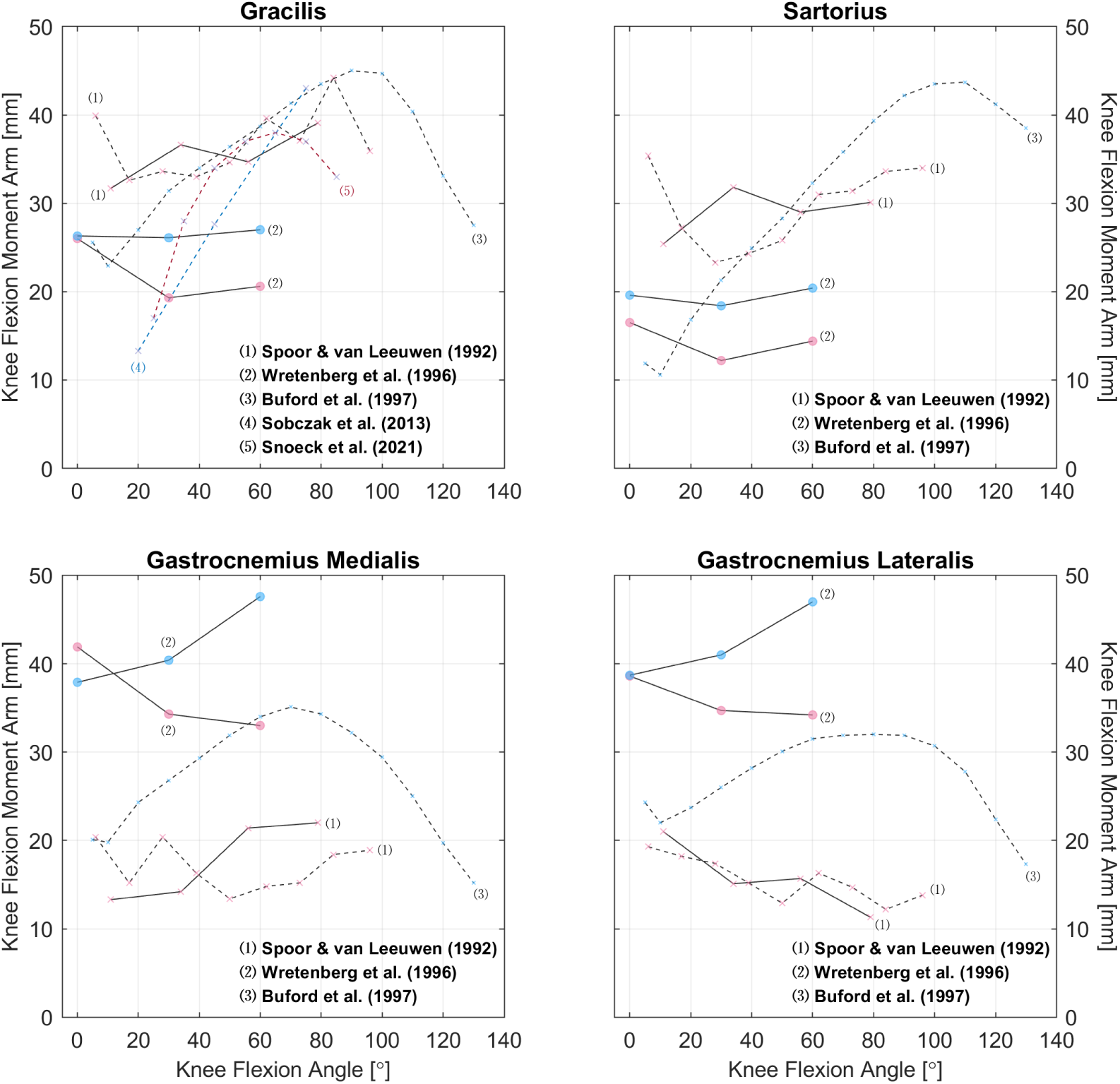
Relations of knee flexion moment arms of other knee flexors. Measurements were performed on healthy subjects (circle) or cadeveric specimens (cross mark). The scatter colors of red, blue, and purple denote respectively the sex of male, female, and mixed/unknown. The solid and dashed line styles denote respectively the CoR and TE methods; the different line and text colors are merely for convenience of distinction.

**Figure 7.**
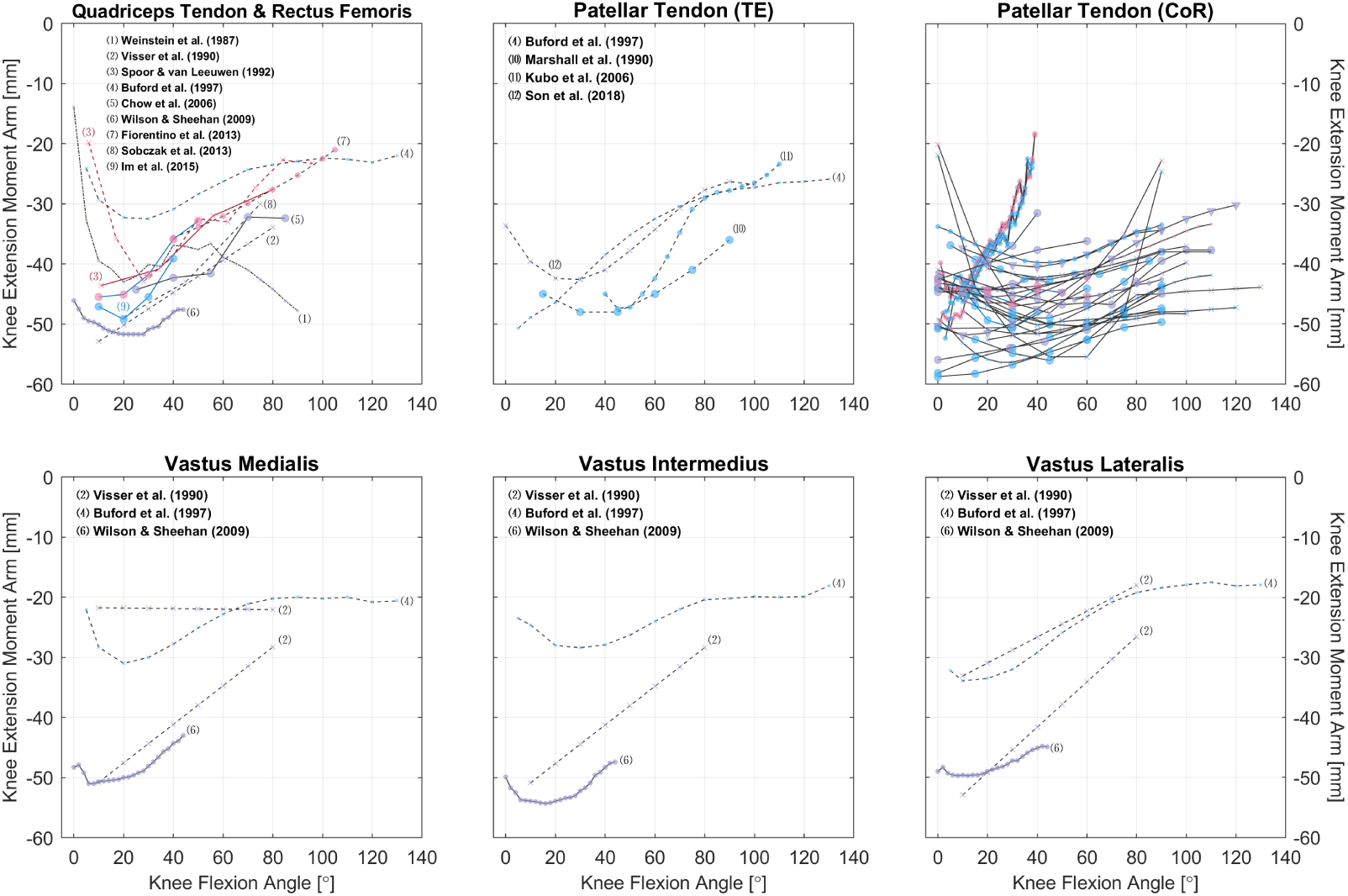
Relations of knee extension moment arms of the quadriceps and their tendons. Measurements were performed on healthy subjects (circle) or cadeveric specimens (cross mark). The scatter colors of red, blue, and purple denote respectively the sex of male, female, and mixed/unknown. The solid, dashed, and dotted line styles denote respectively the CoR, TE, and KE methods; the different line and text colors are merely for convenience of distinction. See Appendix B for detailed references for the patellar tendon.

**Figure 8.**
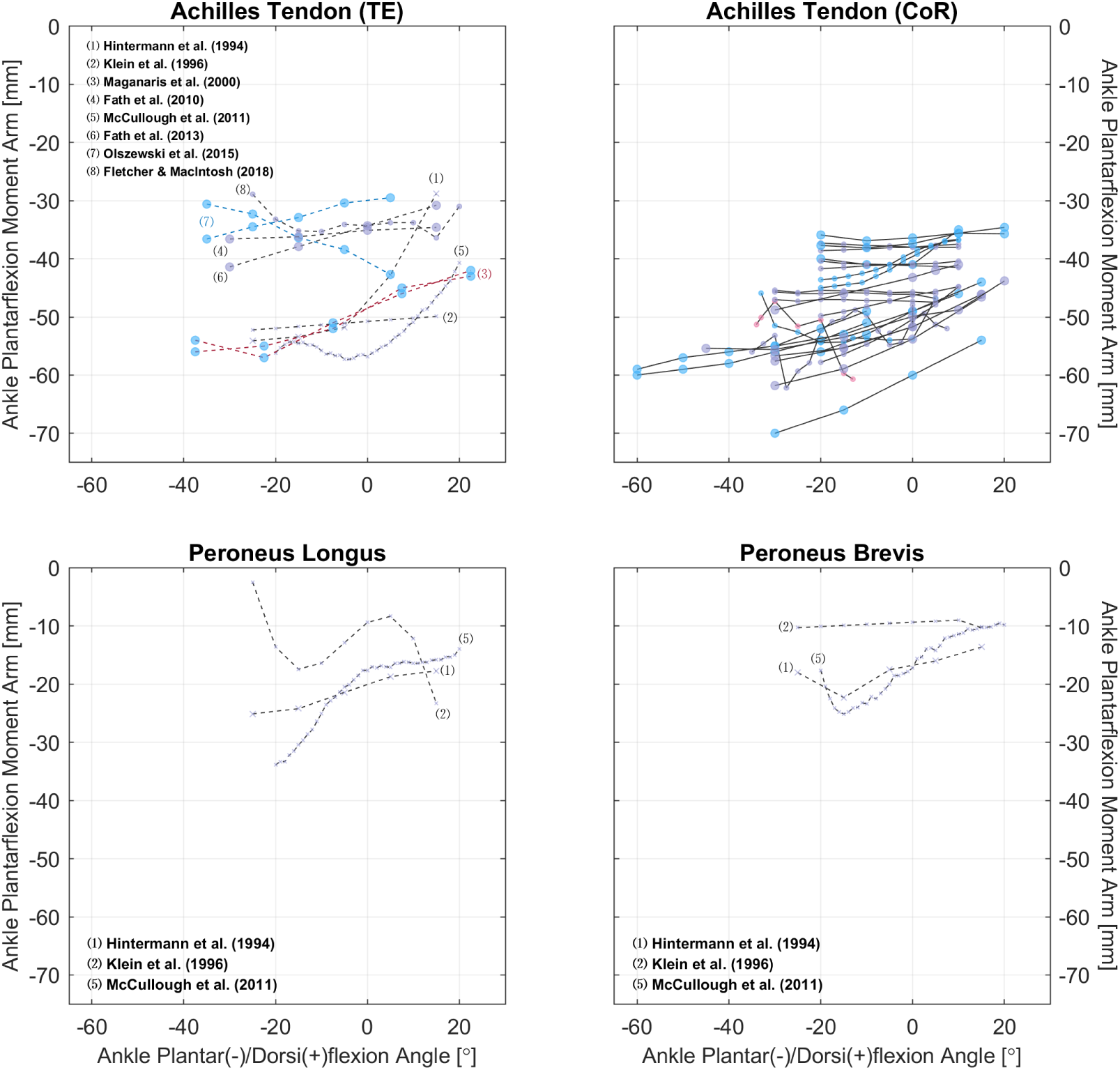
Relations of ankle plantar-/dorsiflexion moment arms of the Achilles tendon and peroneal muscles. Measurements were performed on healthy subjects (circle) or cadeveric specimens (cross mark). The scatter colors of red, blue, and purple denote respectively the sex of male, female, and mixed/unknown. The solid and dashed line styles denote respectively the CoR and TE methods; the different line and text colors are merely for convenience of distinction. See Appendix B for detailed references for the Achilles tendon.

**Figure 9.**
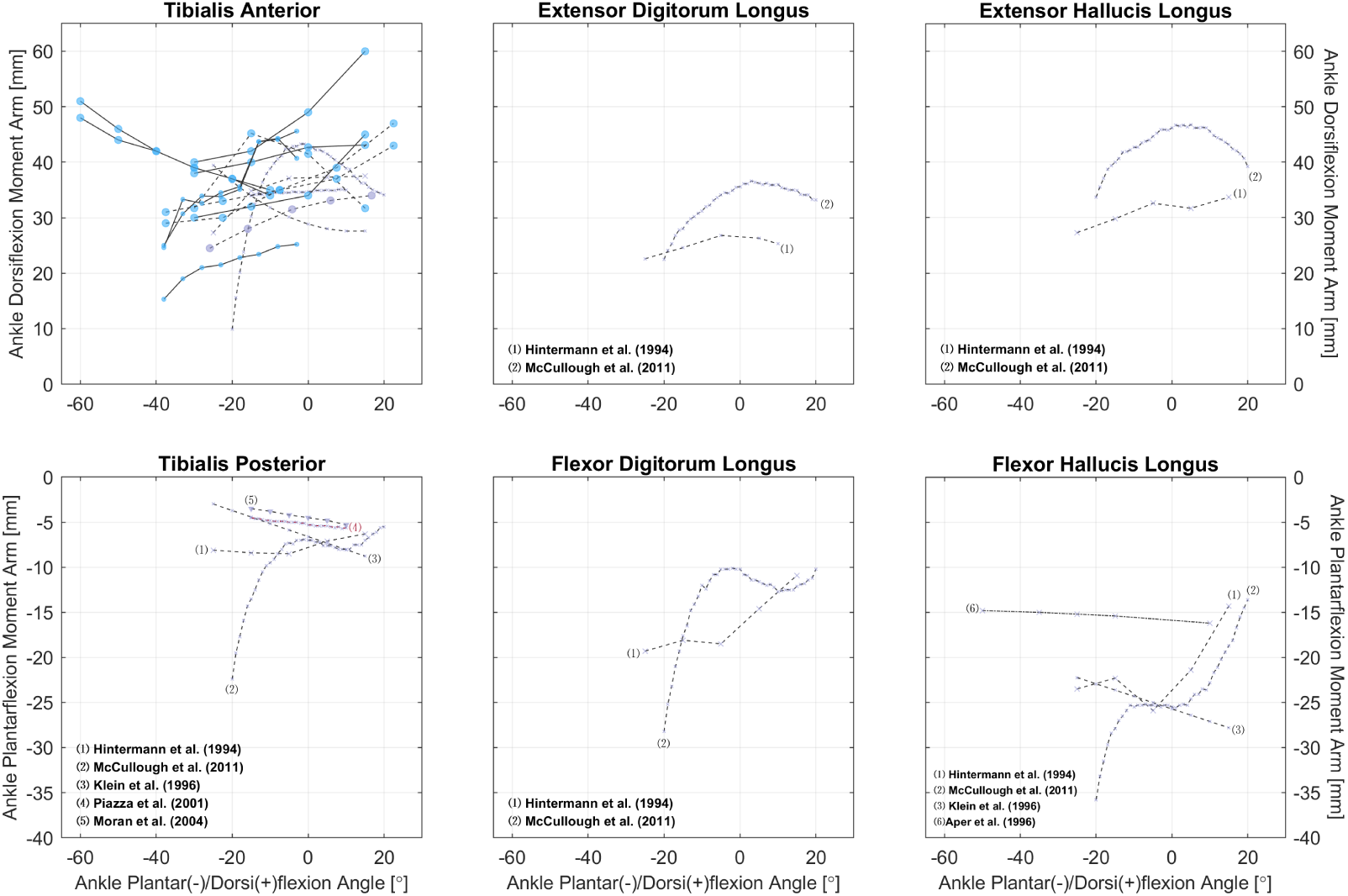
Relations of ankle plantar-/dorsiflexion moment arms of other leg muscles. Measurements were performed on healthy subjects (circle) or cadeveric specimens (cross mark). The scatter colors of blue and purple denote respectively the sex of male and mixed/unknown. The solid, dashed, and dotted line styles denote respectively the CoR, TE, and KE methods; the different line and text colors are merely for convenience of distinction. See Appendix B for detailed references for the tibialis anterior.

**Figure 10.**
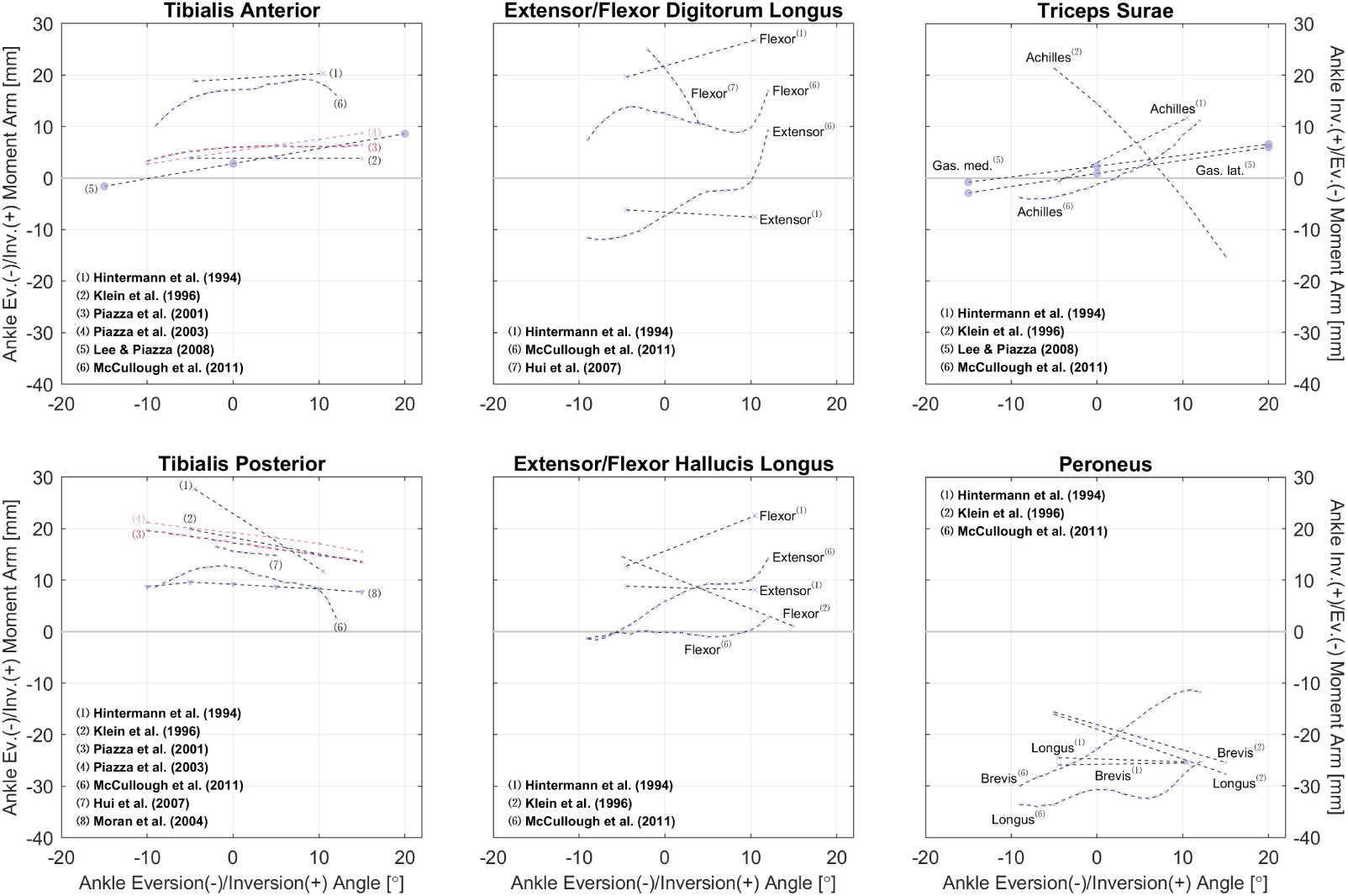
Relations of ankle eversion/inversion moment arms. All measurements were performed on healthy subjects of mixed sex (purple circle) or cadeveric specimens of mixed/unknown sex (purple cross mark) using the TE method (dashed line style); the different line and notation colors are merely for convenience of distinction.

## 4. Discussion

The goal of this study was to gather moment arm datasets and provide references for biomechanical analysis. Such information is especially critical for estimating muscle forces or activations in a given task. In this process, joint moments estimated from kinematic measurements must be distributed to each muscle and transformed into force or activation, both of which are dependent on moment arm. Critically, the accuracy of a musculoskeletal simulation, as well as the reliability of the drawn conclusions, are capped by the quality of moment arm data used for model calibration.

Although we launched a large-scale record search in the database, only a limited number of related studies were identified. This is not entirely surpris-ing, since moment arm is largely an anatomical property, whose measurement is much more difficult than that of kinetic properties such as joint moment or external force. The two primary methods of moment arm measurement requires either cadeveric specimens or medical imaging, both of which can be expensive and time-consuming.

As shown in Table 3, most datasets are for moment arms in the sagit-tal plane, and this matches our finding in gathered joint moment datasets (*citation*: companion submission). In daily locomotion such as walking and running, the dynamics in the sagittal plane are more dominant compared with in the frontal and transverse planes (Camargo et al., 2021; Fukuchi et al., 2017). It is also typical to assume that the knee mainly functions in the sagittal plane, while the ankle does not rotate much. Hence, research in-terests tend to be focused on the sagittal plane. However, the ab-/adduction and ex-/internal rotation of the hip are too evident to ignore for accurate dynamic analysis, yet there are only three studies reporting the moment arm relations in a few related muscles (Figs 3 and 4). For simulations investi-gating the dynamics in non-sagittal planes, it can be difficult to calibrate a musculoskeletal model based on existing datasets, and cautions should be given if conclusions are drawn without specifying the source of calibration data.

Even in the sagittal plane, many muscles lack sufficient data to reveal the characteristics of their moment arm–joint angle relations. In the hip, there are only six studies with relevant data: none is reported for the gluteus minimus, adductor longus/brevis/magnus, pectineus, or gemelli, while for the other hip muscles, there is not a second study with available measurements (Fig 2). Moreover, most ankle muscles lack moment arm data even in the sagittal plane (Figs 8 and 9). Similarly, musculoskeletal simulations may be questionable if they are designed to estimate the activities of these muscles.

Meanwhile, major muscle groups such as the quadriceps femoris, hamstrings and triceps surae are frequently studied with many moment arm measurements. Nevertheless, it is important to note that muscle forces (or activations) are correlated throughout the lower limb: Overestimating the force of a muscle will certainly result in the agonist forces being underestimated and the antagonist forces being overestimated. This means that if too many muscles are not accurately calibrated, even the results of accurately calibrated muscles will likely suffer from error.

Another noteworthy point is that, although the quadriceps femoris, the biceps femoris, and the triceps surae each have its common insertion tendon, some studies report the separate moment arms of individual agonists (Buford et al., 1997; Lee and Piazza, 2008; Wilson and Sheehan, 2009; Visser et al., 1990). This can be confusing because if the muscle line of action is set as the direction of the tendon, then according to the CoR method, muscles with a common tendon share the line of action and should have the same moment arm. However, based on the TE method, or the CoR method with differently defined lines of action, muscles with a common insertion may still have different length variations. For example, to measure excursion, Buford et al. (1997) sutured four cables for each of the quadriceps, and the insertions were selected as “the mid point of each muscle insertion in the quadriceps tendon.” This way, the insertions become independent, and tendon excursion can be different as the joint rotates. Similarly, for each of the quadriceps, Wilson and Sheehan (2009) determined the lines of action as the direction of the muscles, and the measurement using the CoR method naturally differs.

To answer the question of which method is more accurate, it becomes critical to revisit the definition of moment arm as well as tendon anatomy. Essentially, it is implied in the CoR method that if muscles share a common tendon, force is uniformly distributed across the tendon, so the line of action remains unchanged regardless of which muscles are activated. On the other hand, the TE method assumes an independent line of action for each muscle even if they merge into one tendon. Anatomical experiments seem to support the latter. Grob et al. (2016) show that the quadriceps tendon is consisted of three layers, and the components of the second layer (the medial vastus medialis and the vastus lateralis) have different orientations. Mahan et al. (2020) show that the Achilles tendon is a confluence of the gastrocnemii and the soleus, and it can be unbraided into three sub-tendons, and each inserts into their own calcaneal facets. However, despite the anatomical distinction, the sub-tendons are intertwined with transverse interaction, so the force of a muscle might not be entirely transmitted via its own sub-tendon.

In theory, the method of kinetic balance should be the most accurate, where moment arm is calculated from the force measured on the muscle (or its substitute cable) to maintain a constant joint moment. The main advan-tage of this method is based on the kinetic definition of moment arm (Eq 1). Regardless of how muscle force distributes in the tendon, it is directly matched with joint moment: either *in vivo* or *in silico*, the muscle has to generate the same amount of force to exert the same joint moment. In other words, this kinetic-based method eliminates anatomical factors such as ten-don architecture; neither is there the need to consider muscle architecture, tendon curvature, or center of rotation, which is a main concern in the CoR method. A major disadvantage is the inconvenience of measurement, as it requires cadeveric specimens, force transducers, and a special apparatus to apply constant loads. In the studies we collected, only two designed ex-periments are based on the concept of kinetic balance (Aper et al., 1996; Weinstein et al., 1987).

In addition to concerns regarding common tendons, there are pros and cons for both the CoR and TE methods. Generally, the TE method is similar to the KB method, except that it is more convenient. Only length is measured and the experiment can be performed without applying a constant joint load. Most importantly, there is no need to identify the rotation center. This perhaps explains a recent rise of studies performing the TE method *in vivo* with hand-held ultrasound (Fath et al., 2010, 2013; Manal et al., 2013; Miller et al., 2015). A major limitation of this method is that, moment arm is the derivative of of length change with respect to joint angle, so the error in length measurement is easily magnified in derivation. If moment arm is calculated as the quotient of length difference and angle difference, it is typical to see moment arm that zig-zags as joint angle changes (Spoor and van Leeuwen, 1992). If the length data are fitted and moment arm is taken as the derivative of the fitting formula, then its relation with joint angle heavily depends on the choice of formula; e.g., it is linear if the fitting formula is quadratic (Lee and Piazza, 2008; Visser et al., 1990), or quadratic if the formula is cubic (Klein et al., 1996; Vaarbakken et al., 2014, 2015). On the hand, the CoR method directly measures some distance as moment arm, so it is free of this particular problem. However, whether this distance is an appropriate representation of moment arm depends on if the path of the muscle–tendon unit is straight, if the center of rotation is correctly identified, and if the muscle line of action is correctly identified. Due to these restrictions, the CoR method is mostly performed on muscles with large tendons, such as the quadriceps and triceps surae. Nonetheless, the identification of the center of rotation remains a problem for this method. For this, either the Reuleaux method is employed, or some anatomical landmark is assumed as the center. The latter is prone to error (Tsaopoulos et al., 2009; Wade et al., 2019), and if the center of rotation is moving rather than fixed, the Reuleaux method must be frequently repeated to ensure accuracy (Rugg et al., 1990).

Critically, the center of rotation is often estimated in 2D, e.g., the sagittal plane, and the accordant 2D moment arm is in fact a projection of the true 3D moment arm onto the respective plane. In essence, the fundamental motions of the joint, such as hip abduction, knee flexion, or ankle inversion, are not completely confined to the sagittal, frontal, or transverse planes, hence their rotational axes are not strictly perpendicular to these planes. For example, knee flexion, as the motion observed *in vivo*, is mainly rotation in the sagittal plane, but accompanied with some level of rotation in both frontal and transverse planes. A good example is found in Wretenberg et al. (1996), who measured knee moment arms in both sagittal and frontal planes; e.g., the sagittal and frontal moment arms of the patellar tendon are approximately 45 and 5 mm. This does not necessarily mean that the moment actuating knee extension is nine times than the moment actuating knee abduction, but rather knee extension/flexion itself may contain partial rotation in the frontal plane, and the force exerted by the quadriceps could all be transformed into knee extension moment; this case, the true knee extension moment arm of the patellar tendon is 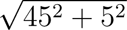 = 45.3 mm. This means that when the moment arm data for calibration are obtained using the 2D CoR method, it is important to check if the rotation axis in measurement matches with that in the musculoskeletal model. If the model rotation axis is not perpendicular to the reference planes, then data obtained via the 2D CoR method will underestimate the model moment arm. Similarly, moment arm measured using the other methods is based on a *natural* rotation axis, so it is important to configure the model with a similar axis. Otherwise if a perpendicular axis is configured, the model moment arm might be overestimated.

Finally, an ideal calibration requires the high-dimensional relations be-tween all moment arms of a muscle and all its related joint angles (Chen et al., 2024). This was one other motive of our study, but most of the gath-ered datasets are about the relation between moment arm and angle in the same plane; except for Hintermann et al. (1994), Vaarbakken et al. (2014), and Wolfram et al. (2018). For accurate musculoskeletal modeling, it is nec-essary to obtain moment arm data for as many actuating DoFs in as many muscles, and Table 3 should serve as a good reference to experiment design for gap-filling measurements. Currently, several approaches may be taken in compensation for the lack of experimental data. If 3D MRI reconstruction is available, moment arm can be estimated using the CoR method. For conve-nience, only one position is needed, and the rotation axes for measurement can be matched with those configured in the musculoskeletal model. This way, a rough estimation of the moment arm magnitude can be obtained. As a less preferable alternative, one could model an initial muscle path based on the anatomical origin and insertion points in the skeletal model, and take the distance between muscle path and rotation axis as moment arm. With the estimated magnitude, one must then assume a certain relation for the variation of moment arm in different joint positions. This could be assumed as constant, or similar to that of the agonists. For example, the gluteus maximus and medius tend to function as hip external rotators when the hip is extended or slightly flexed, and they gradually become internal rotators as in highly flexed hip positions (Delp et al., 1999). So a similar pattern can be assigned to the gluteus minimus. In contrast, for hip muscles that are not typically known as rotators, their rotation moment arms can be treated as a small constant. As a last step, with the estimated magnitude and as-sumed relations, moment arms may be generated for various joint positions and serve as calibration data for automated muscle path calibration (Chen et al., 2024).

## 5. Conclusion

A total of 300 moment arm datasets were collected from literature to illustrate muscle moment arm–joint angle relations in the hip, knee, and ankle. The overall pattern and magnitude are presented for six actuated DoFs. The findings should provide valuable insight into musculoskeletal mechanics, enhance biomechanical analysis, and improve the design of future experiments for moment arm measurement.

## Acknowledgements

This work was supported by the Lighthouse Initiative Geriatronics by StMWi Bayern (Project X, grant no. 5140951).

## Appendix A. Supplementary material

Files associated with this article, including the search log and metadata catalog (.xlsx), datasets (.mat), and visualization scripts (.m), can be found online at *link-to-add*.

## Appendix B. Summary of studies and datasets

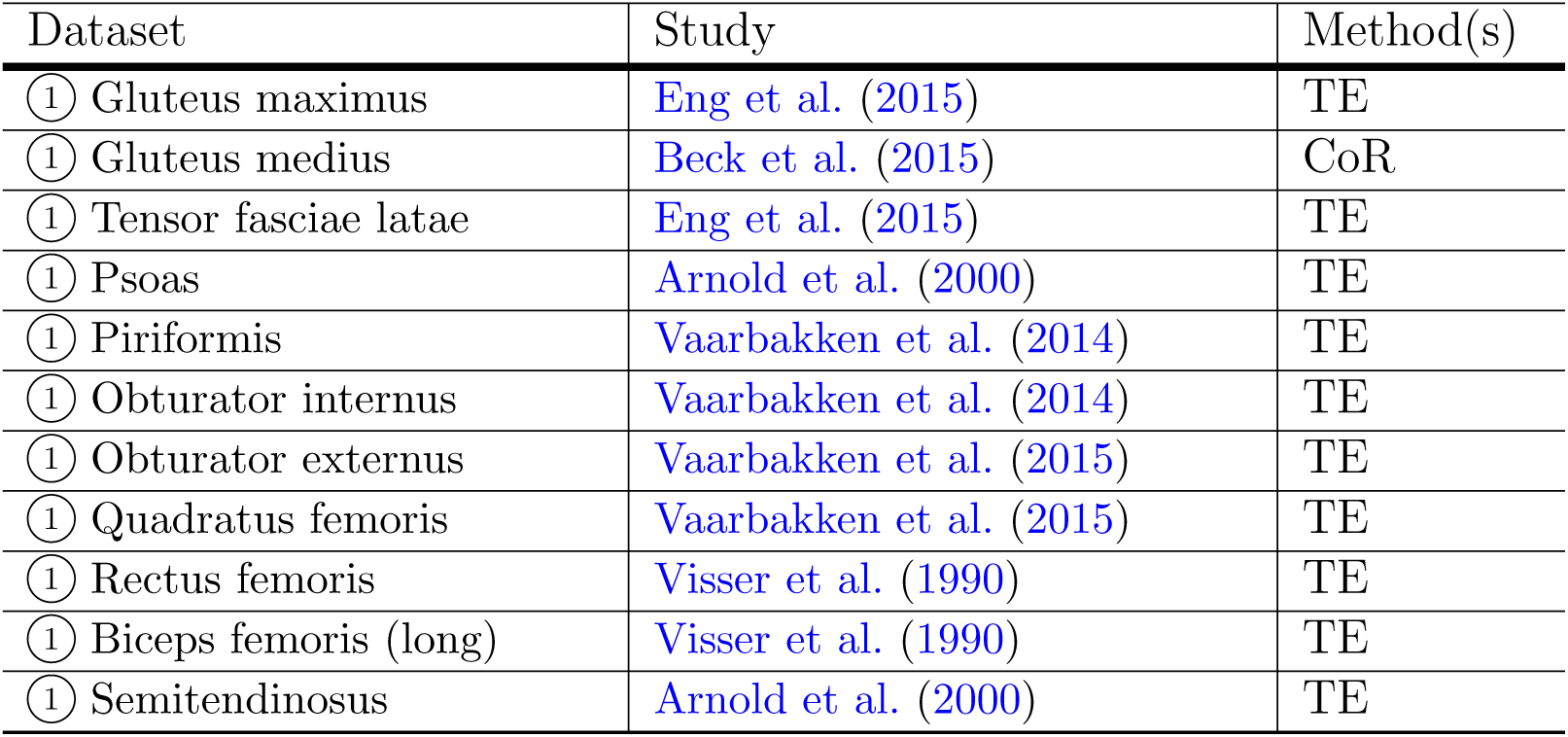
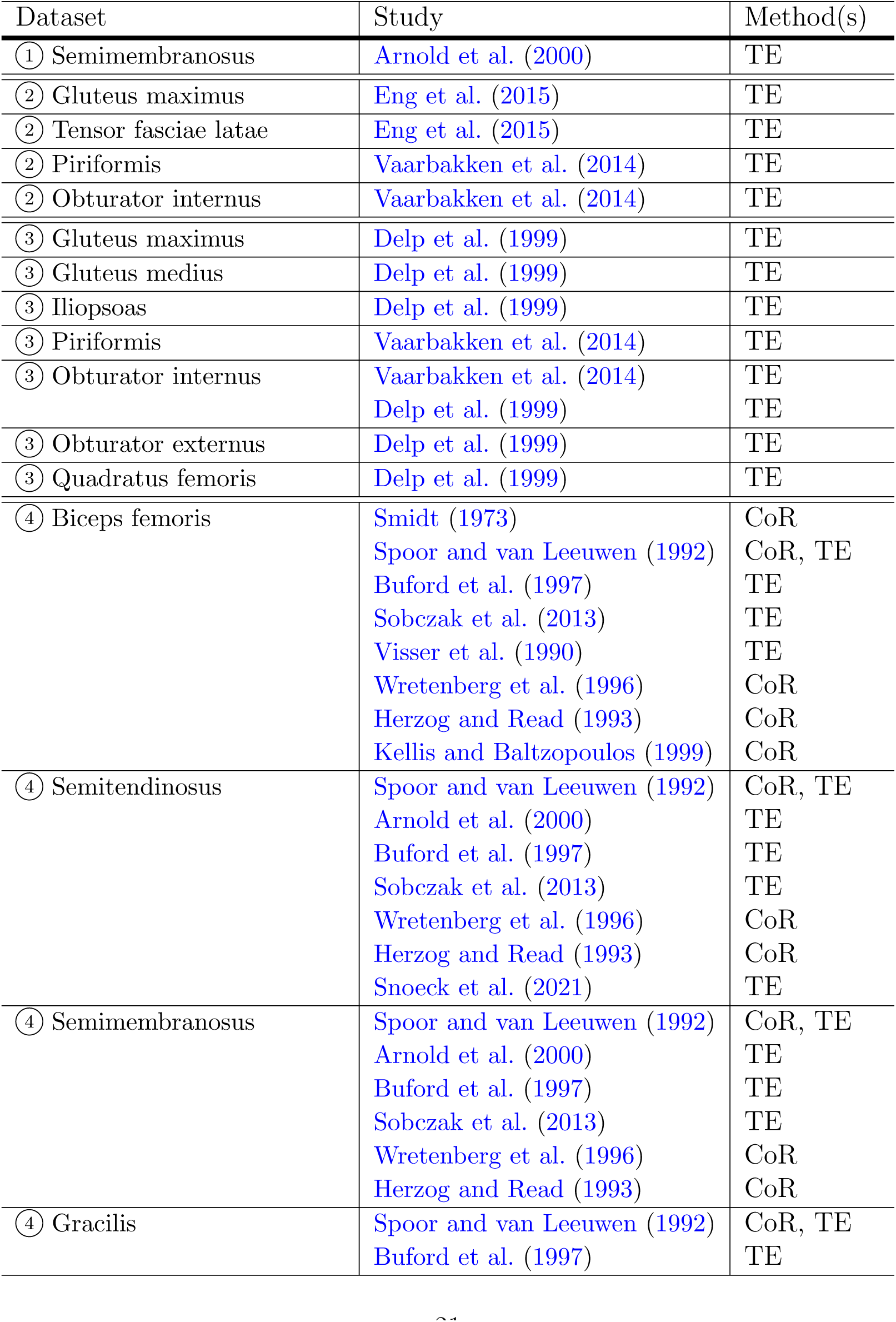
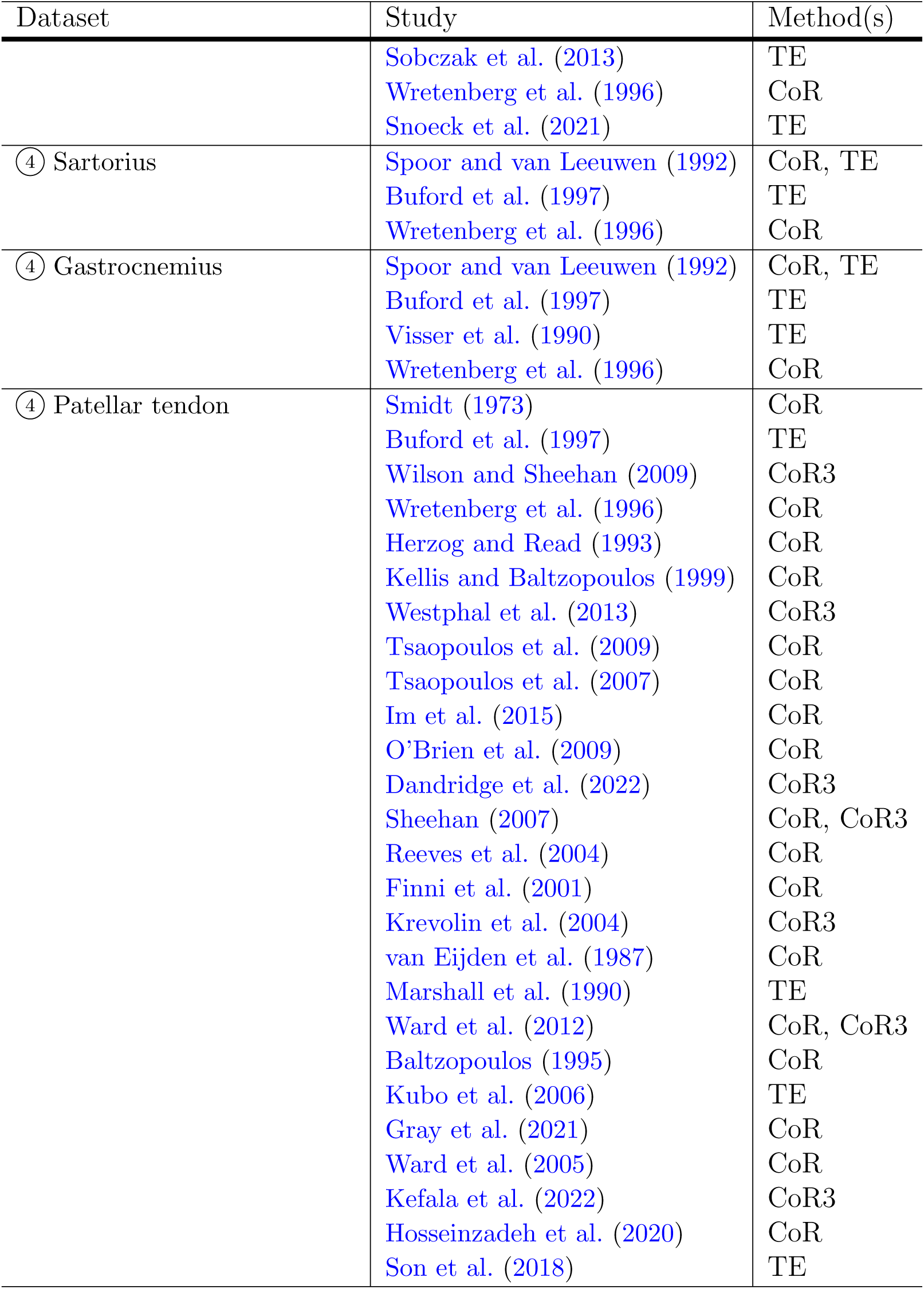
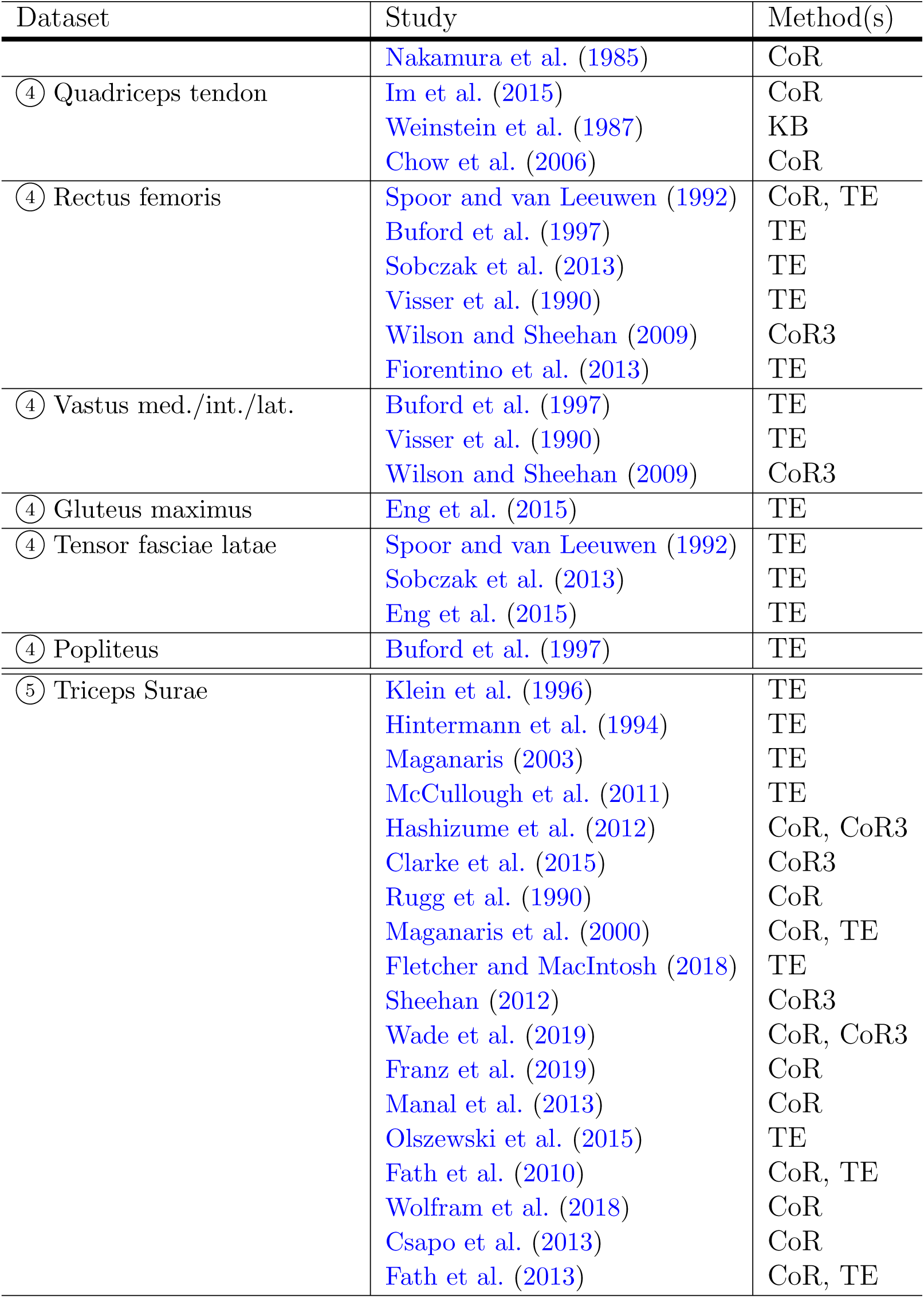
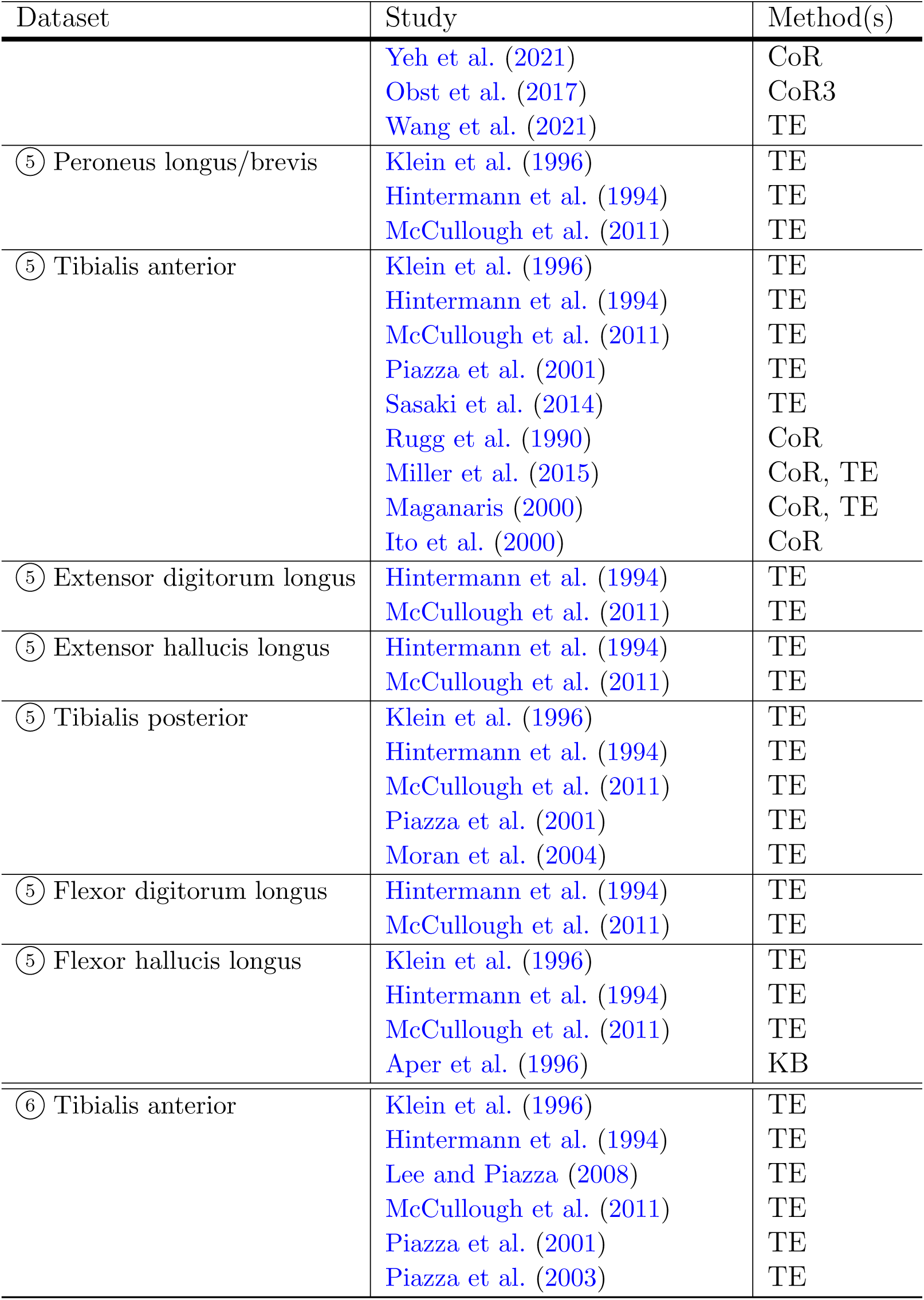
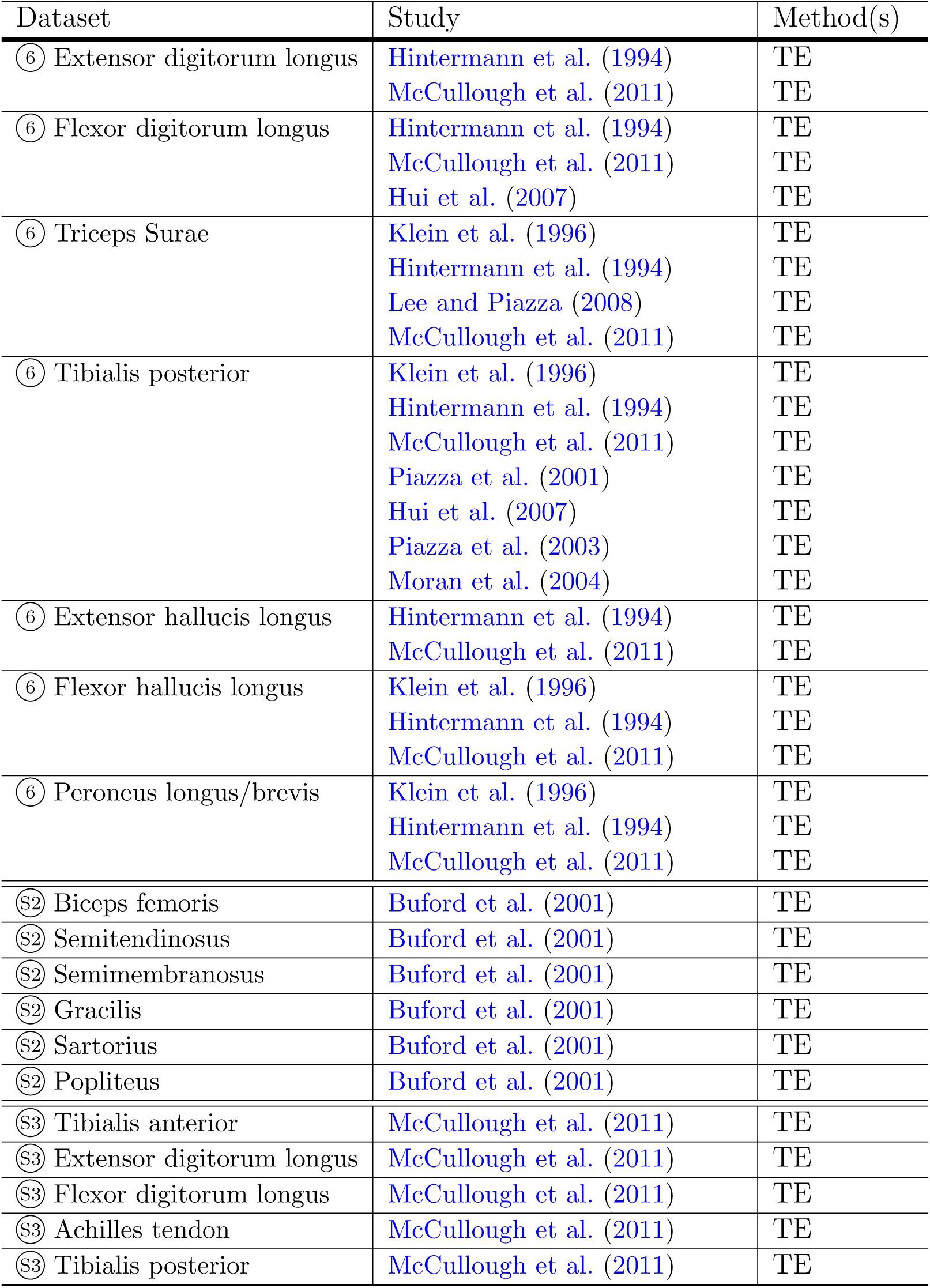
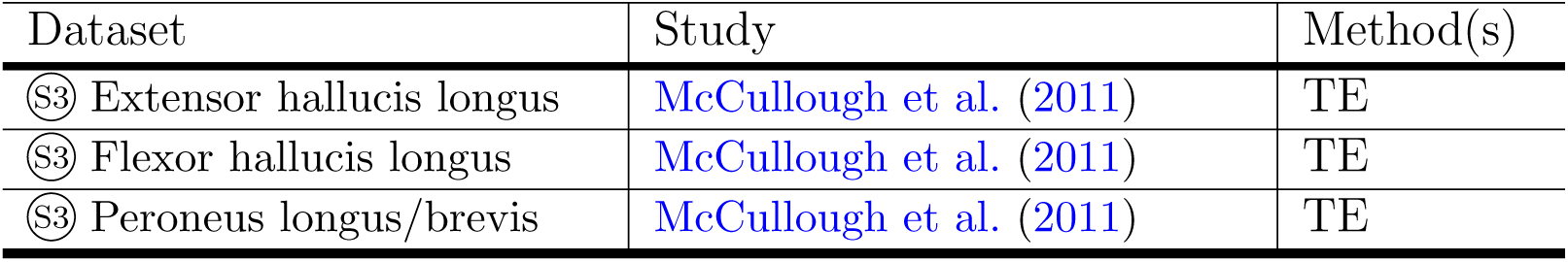

